# Neural Mechanisms for Executive Control of Speed-Accuracy Tradeoff

**DOI:** 10.1101/773549

**Authors:** Thomas R. Reppert, Richard P. Heitz, Jeffrey D. Schall

**Affiliations:** Center for Integrative & Cognitive Neuroscience, Vanderbilt Vision Research Center, Department of Psychology, Vanderbilt University, Nashville, TN 37240, USA

**Keywords:** actor-critic, cognitive control, supplementary eye field, medial frontal cortex, performance monitoring, reward prediction error

## Abstract

The balance of speed with accuracy requires error detection and performance adaptation. To date, neural concomitants of these processes have been investigated only with noninvasive measures. To provide the first neurophysiological description, macaque monkeys performed visual search under cued speed accuracy tradeoff (SAT). Monkeys changed SAT emphasis immediately after a cued switch while neuron discharges were sampled in medial frontal cortex area supplementary eye field (SEF). A multiplicity of SEF neurons signaled production of choice errors and timing errors. Modulation of SEF activity after choice errors predicted production of un-rewarded corrective saccades. Modulation of activity after timing errors signaled reward prediction error. Adaptation of performance during SAT of visual search was accomplished through pronounced changes in neural state from before search array presentation until after reward delivery. These results contextualize previous findings using noninvasive measures, complement neurophysiological findings in visuomotor structures, endorse the role of medial frontal cortex as a critic relative to the actor instantiated in visuomotor structures, and extend our understanding of the distributed neural mechanisms of SAT.

**HIGHLIGHTS:**

- Medial frontal cortex enables post-error adjustment during SAT
- Choice and timing errors were signaled by partially overlapping neural pools
- Medial frontal cortex can proactively modulate visuomotor processes
- Medial frontal cortex is to visuomotor circuits as critic to actor

## INTRODUCTION

The speed-accuracy tradeoff (SAT) is a classic paradigm for investigating mechanisms of decision making (Heitz, 2014). Human studies of SAT with noninvasive measures have offered insights but highlight uncertainty about fundamental questions.

Multiple event-related potential (ERP) studies of SAT have been conducted (van der Lubbe et al., 2001; Osman et al., 2000; Rinkenauer et al., 2004; Sangals et al., 2002; Wenzlaff et al., 2011). In general, event-related potential evidence suggests SAT operates on attention allocation and response preparation. Other ERP studies have investigated error monitoring when Fast or Accurate responses are cued. Previous studies reported that emphasis on accuracy enhanced the ERN and Pe (e.g., Arbel and Donchin, 2009; Falkenstein et al., 2000; Gehring et al., 1993). However, other investigators found the opposite (Steinhauser and Yeung, 2012). Such a fundamental disagreement can arise for multiple reasons, but clarity is gained through neurophysiological investigation of one source contributing to the ERN and Pe, the SEF (Sajad et al. 2019).

Multiple fMRI studies of SAT have been conducted (Forstmann et al., 2008; Ivanoff et al., 2008; van Maanen et al., 2011; van Veen et al., 2008; Weigard et al., 2018). Different tasks and analysis pipelines lead to different cortical and subcortical regions identified or emphasized across studies. However, all fMRI investigations of SAT feature medial frontal areas, in particular preSMA. Some studies report greater activation of preSMA with an emphasis on speed during the baseline period (Forstmann et al., 2008; van Veen et al., 2008). Other studies report a similar effect during the response time interval (Ivanoff et al., 2008; van Maanen et al., 2011; van Veen et al., 2008; Weigard et al., 2018). Several investigators propose that preSMA sets the excursion between baseline and threshold levels of an evidence accumulation process through interactions with striatum and other regions (Forstmann et al., 2008; Ivanoff et al., 2008; van Maanen et al., 2011; Weigard et al., 2018; Wenzlaff et al., 2011). This proposal entails assumptions about the neural processes in areas like preSMA that can only be evaluated with neurophysiological measures. We evaluated these assumptions with a neurophysiological investigation of the oculo-motor counterpart of the skeletal-motor preSMA, the SEF.

The SEF, located on the dorsal convexity of medial frontal cortex in macaques, contributes indirectly to saccade production (e.g., Stuphorn et al., 2010), just as preSMA contributes indirectly to limb movements (e.g., Scangos and Stuphorn, 2010). Also, in difficult tasks when gaze errors can occur, performance monitoring signals are evident in SEF (Abzug and Sommer, 2018; Emeric et al., 2010; Kawaguchi et al., 2015; Sajad et al., 2019; So and Stuphorn, 2012; Stuphorn et al., 2000), just as such signals are evident in preSMA (e.g., Isoda and Hikosaka, 2007; Scangos et al., 2013). These common functions are paralleled by common patterns of anatomical connectivity of SEF (e.g., Huerta and Kaas, 1990) and preSMA (e.g., Luppino et al., 1993), including to striatum (Parthasarathy et al., 1992).

Here we describe the first neurophysiological recordings obtained from medial frontal cortex of macaque monkeys performing a perceptual decision-making task under SAT. We relate the findings to previous investigations of SEF during different tasks that offer insight on the assumptions made by non-invasive studies (e.g., Stuphorn et al., 2000, 2010) and to previous neurophysiological investigations of SAT in key nodes in visuo-motor networks (Heitz and Schall, 2012; Reppert et al., 2018) (see also Hanks et al., 2014; Thura and Cisek, 2016). All of these findings can be interpreted through neurocomputational modeling of SAT based on the data from FEF (Servant et al., 2019). The mechanistic view emerging from all of these new results complements, contradicts, and extends the current, most commonly accepted view.

## RESULTS

### Speed-accuracy tradeoff of performance

Two macaque monkeys performed a visual search task to locate a target item presented amongst seven distractor items (Figures 1A, 1B). Trials began when monkeys fixated a central stimulus, the color of which cued emphasis on speed (Fast condition) or accuracy (Accurate condition) of responding. Search contingency was fixed as either T amongst L’s (more efficient or less difficult) or L amongst T’s (less efficient or more difficult). Monkeys were trained extensively on the mapping between cue color and task condition. Task condition alternated every 5 or 10 trials. Reward (fluid) and punishment (time-out) contingencies were employed to shape and maintain the SAT. Response deadlines (Fast: 392 ± 17 ms, Accurate: 443 ± 6 ms) were similar to those employed in human studies (Heitz and Engle, 2007; Rinkenauer et al., 2004; Wickelgren, 1977).

**Figure 1.**
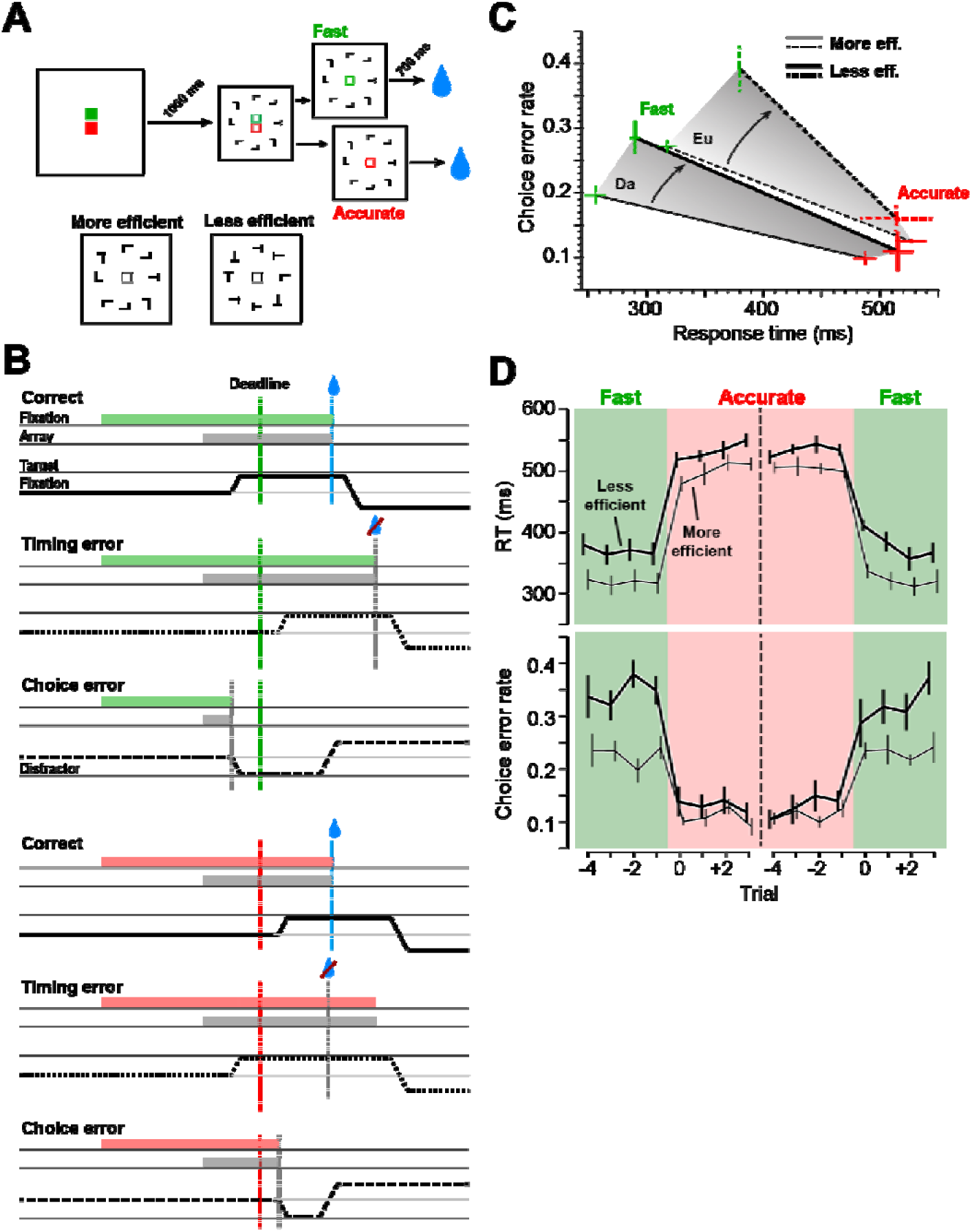
Speed-accuracy tradeoff of visual search. (**A**) Trials began with fixation of a central stimulus that cued Fast (green) or Accurate (red) responding, after which an iso-eccentric array of 8 T/L form shapes appeared. Monkeys searched for the target item (rotated T or L) presented amongst seven distractors (rotated L or T). Fluid reward was delivered after 700 ms of fixation of the target item. Fast and Accurate trials alternated in short blocks. Monkeys were more efficient (faster RT, fewer errors) performing search for T among L and less efficient performing search for L among T. (**B**) Timeline of events preceding trial outcomes in the Fast (above) and Accurate (below) conditions. On correct trials, monkeys shifted gaze to the target with appropriate timing (solid black lines). Monkeys occasionally committed timing errors (dotted lines) or choice errors (dashed lines). In the Accurate and Fast conditions, timing errors were committed before and after the deadline, respectively. (**C**) Tradeoff of RT and choice error rate. RT was shorter, and choice error rate, higher, in the Fast relative to the Accurate condition. Tradeoff curves shifted to poorer performance during less efficient (more difficult) relative to more efficient (less difficult) search. Error bars are SE across sessions. (**D**) Changes in mean ± SE RT (above) and choice error rate (below) were immediate after SAT cue changes. Data are plotted separately for more efficient (thin lines) and less efficient (thick lines) search sessions.

Under speed constraints, RT was shorter (F(1,24) = 387, p = 2.6 × 10^−16^, BF > 1,000), and choice error rate, higher (F(1,24) = 31.4, p = 9.2 × 10^−6^, BF > 1,000), in the Fast relative to the Accurate condition (Figure 1C). During less efficient (more difficulty) search, the tradeoff of RT and error rate shifted to poorer performance with longer RT (F(1,24) = 5.29, p = 0.03, BF = 2.5) and more errors (F(1,24) = 8.6, p = 0.007, BF = 5.9). We found a significant interaction of SAT condition and search efficiency, with SAT being more pronounced during less efficient (more difficult) search (F(1,24) = 4.6, p = 0.04, BF = 1.7). As described previously (Heitz and Schall, 2012; Reppert et al., 2018), the SAT-related changes in RT and choice error rate were immediate upon a cued change in SAT condition (Figures 1D, S1A, S1B).

In summary, we observed pronounced tradeoff of RT and choice error rate during visual search. The tradeoff curve shifted to poorer performance with a greater effect of SAT during less efficient (more difficult) search, as monkeys committed more errors with slower RT. Changes in behavior were immediate upon a cued change in task condition.

### Choice errors and correction

Previous research described modulation of SEF activity after response errors during diverse tasks, including cued detection (Isoda and Hikosaka, 2007), saccade countermanding (Stuphorn et al., 2000; Sajad et al., 2019), and visual search (Purcell et al., 2012). We observed 28 SEF neurons (14 Da, 14 Eu) that signaled production of choice errors, of which 24 (12 Da, 12 Eu) were enhanced, and 4 (2 Da, 2 Eu) suppressed (Figure 2). This modulation was not dependent on activity possibly associated with generation of the second saccade (Sajad et al., 2019) (Figures S2F, S2G). We focused our analyses on the 24 neurons with error-enhanced signaling.

**Figure 2.**
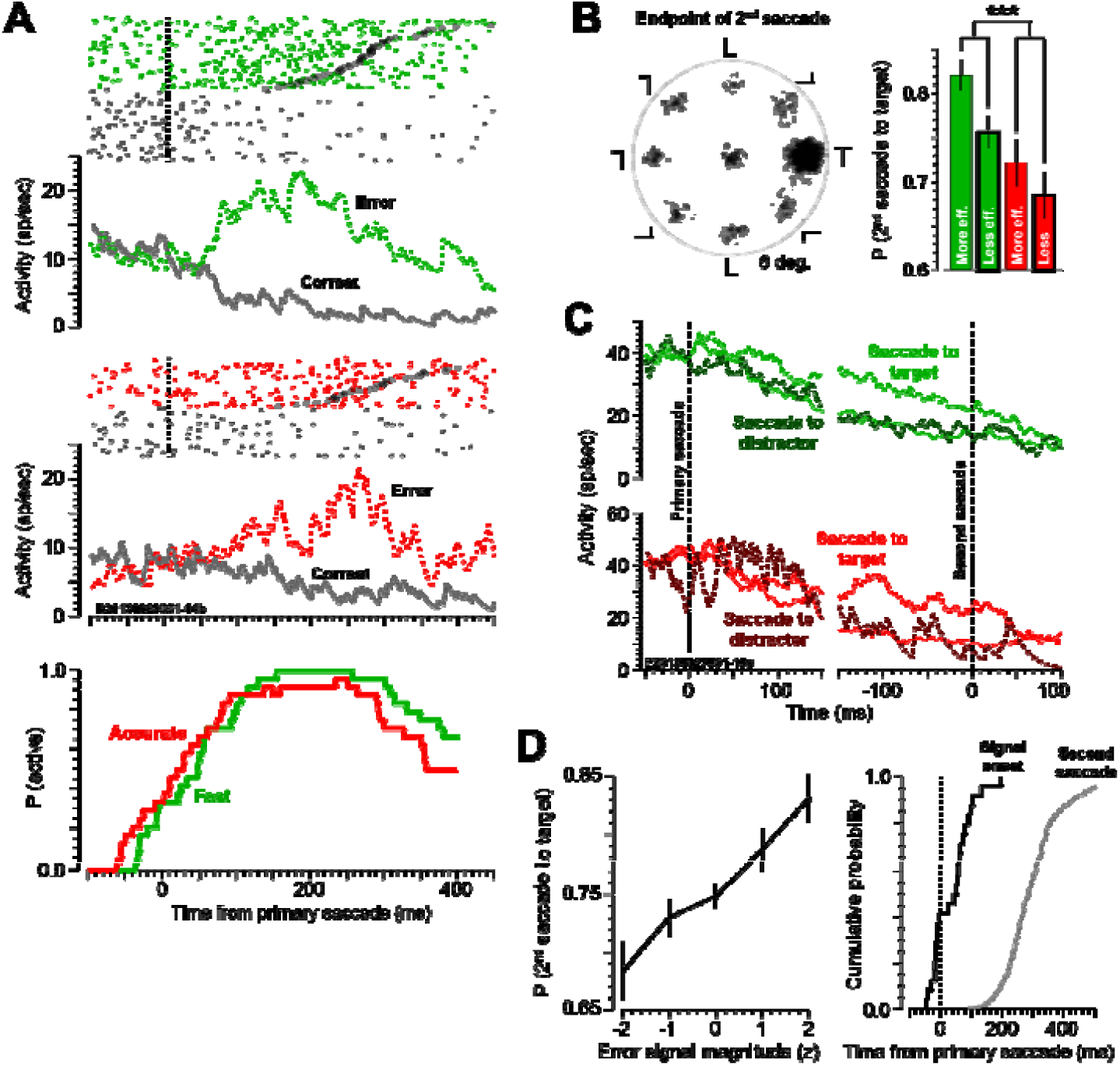
Signaling choice errors and error correction. (**A**) Modulation after choice errors shown with raster and SDF from a representative neuron (above) and probability density of modulation across the sample (beneath). Colored (green/red) and gray rasters represent error and correct trials, respectively. Dark circles in colored rasters are time of the second saccade. The latency and magnitude of the choice error signal was equivalent across task conditions. (**B**) (left) Second saccade endpoint, plotted with target rotated to horizontal right. Data combined across task conditions. (right) Mean ± SE probability of second saccade to target. The likelihood of a corrective saccade to the target was lower during Accurate relative to Fast trials. (**C**) Average SDF for a representative neuron aligned on first saccade (left) and second saccade (right), with second saccade directed to target location (light dashed), or directed to a distractor (dark dashed). The neuron fired more vigorously before corrective saccades, and the modulation ceased after the second saccade. (**D**) (left) Mean ± SE probability of second saccade directed to the target as a function of quantiles of z-scored error-related activity across neurons. Across the population of error-signaling neurons (n = 24), the likelihood of a corrective saccade to the target increased with magnitude of the error signal. (right) Cumulative distributions of time of onset of choice error-related modulation and time of second saccade initiation. Error-related modulation preceded the second saccade by ∼ 200 ms on average. Trials were collapsed across Fast and Accurate task conditions.

Choice error signaling arose ∼ 50 ms before execution of the primary saccade, becoming maximal 100-200 ms thereafter and persisting for up to 500 ms (Figures 2A, S2A). The latency of the choice error signal was equivalent across task conditions (t_23_ = 0.93, p = 0.36, BF = 0.32), as was the magnitude of the signal (t_23_ = 1.69, p = 0.11, BF = 0.73).

After choice errors, monkeys produced a second saccade that was often directed to the location of the target. However, this saccade was directed to the location of a distractor on ∼ 1/4 of trials (Figures 2B, S2B). The likelihood of a corrective saccade to the target was lower during Accurate relative to Fast trials (F(1,24) = 13.89, p = 0.001, BF = 64) and during more efficient relative to less efficient search (F(1,24) = 4.95, p = 0.036, BF = 3.54). The latency of the second saccade was prolonged in the Accurate relative to the Fast condition (Figure S2C) (F(1,24) = 4.30, p = 0.049, BF = 1.72). These new data demonstrate that monkeys encoded the location of the target even though choice errors during visual search arise when neurons in frontal eye field and superior colliculus misidentify a distractor as the target (Reppert et al., 2018) (see also Murthy et al., 2007)).

A relationship between error signals and performance adjustments has been found in some (Gehring et al., 2012; Rodriguez-Fornells et al., 2002; Sajad et al., 2019) but not all studies (Dudschig and Jentzsch, 2009; Fiehler et al., 2005; Gehring and Fencsik, 2001; Hajcak et al., 2003). We found that the likelihood of a corrective saccade to the target increased with magnitude of the error signal (Figures 2D, S2D) (F(4,109) = 9.12, p = 2.2 × 10^−6^). Moreover, the choice error signal started ∼ 200 ms prior to and lasted through the execution of the second saccade (Figure S2E).

In summary, we discovered prominent signaling of choice errors in SEF during SAT of visual search. This signal became maximal ∼ 150-200 ms after generation of the error response and lasted up to ∼ 400 ms. Signal latency and magnitude were equivalent across SAT conditions. After the error saccade, monkeys generated a second saccade, often shifting gaze to the location of the foregone target. This saccade was less often directed to the target in the Accurate relative to the Fast condition. We found that signaling of choice errors in SEF preceded the second saccade by ∼ 200 ms and was related to the endpoint of the second saccade. More vigorous signaling of errors was associated with a higher likelihood of a corrective saccade.

### Timing errors and reward prediction

Previously, we showed that the monkeys’ natural preference was for quick responding, at the expense of accuracy (Heitz and Schall, 2012). As a result, monkeys committed more timing errors in the Accurate relative to the Fast condition (F(1,24) = 51.2, p = 2.15 × 10^−7^, BF > 1,000), indicating a reluctance to withhold responding. Unlike rate of choice errors, rate of timing errors was unaffected by search efficiency (F(1,24) = 0.12, p = 0.74, BF = 0.36).

As noted above, monkeys adapted RT immediately upon a cued change in task condition (Figure 1D). However, they required several trials to fully adjust RT. To test this, we analyzed timing error rate as a function of trial and found that the error rate was greatest on the first trial after a cued condition switch (Figures 3A, S3A) (F(7,104) = 3.5, p = 0.002, BF = 16.7, one-way ANOVA). After the first Accurate trial, monkeys reduced the fraction of short latency saccades produced before the response deadline (Figures 3B, S3B), thus resulting in lower timing error rate. These data suggest that SAT required distinct states of responding in the two task conditions. Transitions between states began immediately after SAT cues changed but multiple trials were needed to complete the adjustment.

**Figure 3.**
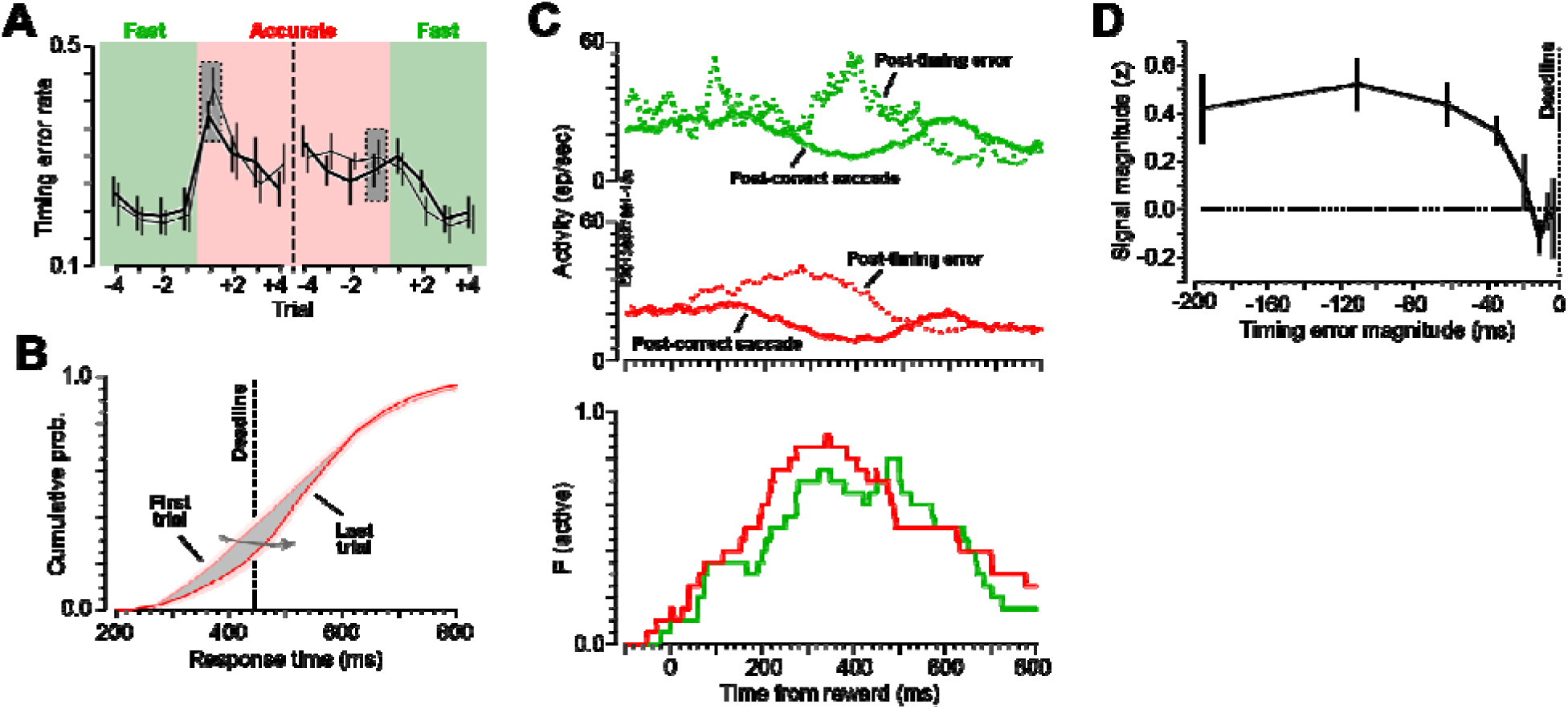
Signaling reward prediction error after timing errors. (**A**) Mean ± SE rate of saccade timing errors relative to cue changes. Changes in timing error rate were immediate upon a cued change in task condition. Gray boxes show trials from which distributions are drawn in (B). (**B**) Cumulative distributions of RT across the first and last trials of the Accurate condition blocks. Most of the RT adaptation was in the earliest responses. (**C**) Modulation of activity after timing errors shown from a representative neuron (above) and probability density of modulation across the sample (beneath) for Fast (green) and Accurate (red) trials. (**D**) Magnitude of z-scored RPE modulation as a function of the magnitude of the timing error relative to the SAT deadline (indicated).

As mentioned, visual feedback after timing errors mirrored feedback on correct trials (Figure 1B). This created an expectation for juice reward after the experienced delay of ∼ 700 ms after the saccade; however, after timing errors, juice reward was withheld. At the time of expected reward, we observed pronounced modulation among particular units. The modulation resembled that described in previous work, described as a reward prediction error (RPE) signal (Amador et al., 2000; Kawaguchi et al., 2015; Nakamura et al., 2005; Sajad et al., 2019; So and Stuphorn, 2012). We found 23 SEF neurons (13 Da, 10 Eu) that signaled RPE at the time of expected delivery. Because the rate of timing errors was lower in the Fast condition, we were unable to estimate the RPE signal for 3 neurons (monkey Eu). Of the remaining 20 neurons, 14 (8 Da, 6 Eu) were facilitated, and 6 (5 Da, 1 Eu), suppressed, at the time of expected reward. The RPE signal arose at the time of expected reward, becoming maximal ∼ 350 ms thereafter and persisting for up to 800 ms (Figures 3C, S3C).

Task condition affected both timing and magnitude of the RPE signal (Figure S3D). The signal arose decisively earlier in the Accurate condition (t_19_ = 4.2, p = 0.0005, BF = 72), but with greater magnitude in the Fast condition (t_19_ = 2.55, p = 0.02, BF = 2.9). We also assessed the relationship between the magnitude of the timing error and the magnitude of the associated error signal. The magnitude of the error signal increased significantly with the magnitude of the timing error (Figures 3D, S3E) (F(7,107) = 4.52, p = 0.0002, one-way ANOVA).

In summary, we assessed the rate of timing errors and an associated RPE signal in SEF. We found that timing error rate was highest on the first trial post-cued condition switch, evidencing a distinct state change between task conditions. After commitment of timing errors, SEF neurons signaled RPE at the time of expected juice delivery, with the signal arising around the time of expected reward and becoming maximal ∼ 350 ms thereafter. The magnitude of the prediction error signal was related to the magnitude of the behavioral error, with more egregious timing errors signaled more vigorously.

Subpopulations of SEF neurons signaled both choice errors and timing errors (n = 8; 3 Da, 5 Eu), while others signaled only choice errors (n = 10; 4 Da, 6 Eu), and others signaled only timing errors (n = 12; 10 Da, 2 Eu) (Figure 4A). This variation was on a continuum. We quantified this (Figure 4B) by calculating contrast ratios for the responses in errors relative to correct trials *[(A*_*error*_ *– A*_*correct*_*)/(A*_*error*_ *+ A*_*correct*_)] where for choice errors *A*_*error*_ and *A*_*correct*_ are the average SDF values 100-300 ms after median time of choice error (Fast condition), and for timing errors *A*_*error*_ and *A*_*correct*_ are the average SDF values 100-500 ms after median time of expected reward (Accurate condition).

**Figure 4.**
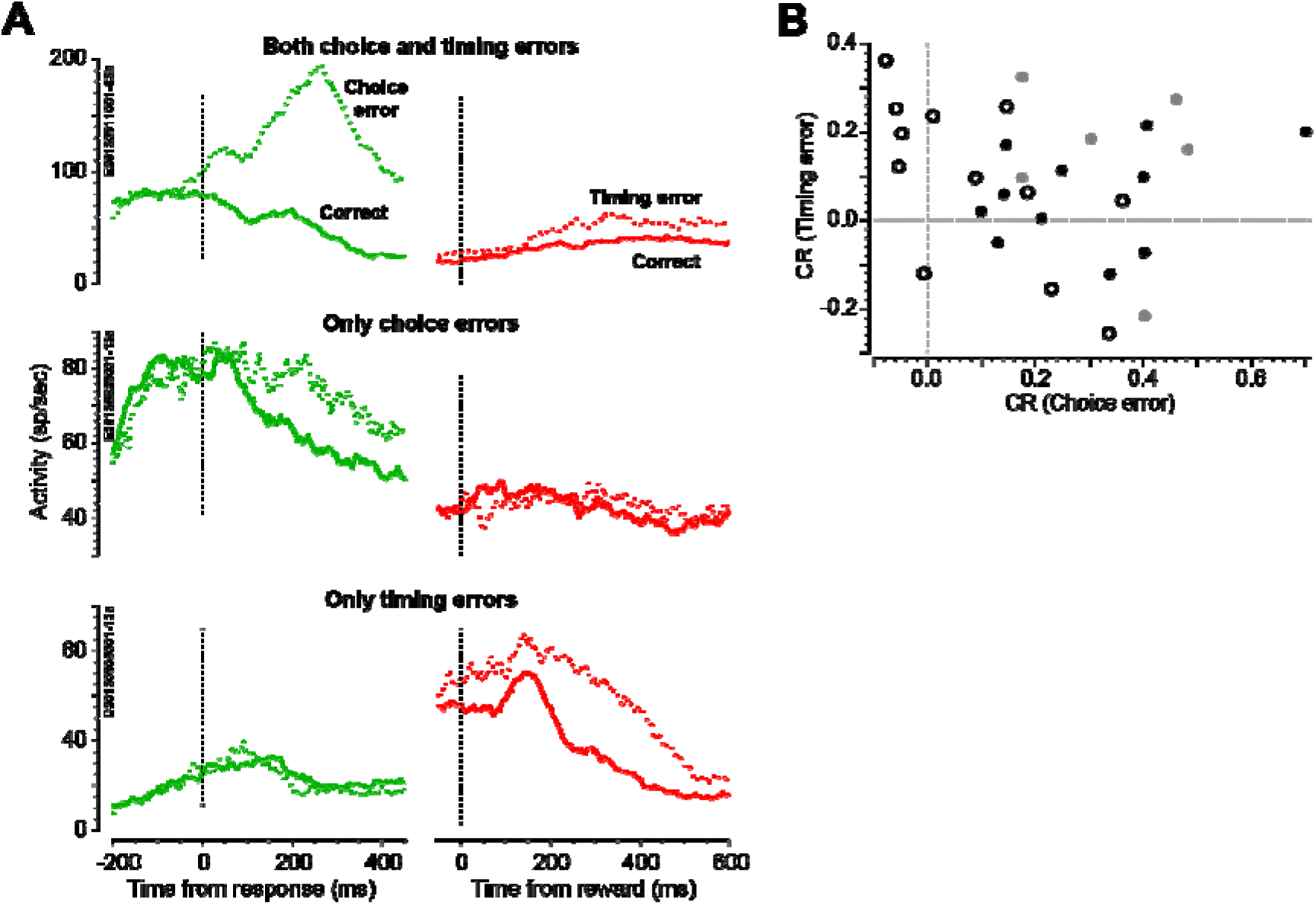
Distinct neurons signaling choice error and reward prediction error. (**A**) Representative neurons signaling both choice error and timing error (top), choice error only (middle), and timing error only (bottom). (**B**) Plot of the relationship between contrast ratios for choice errors and timing errors. Each dot represents a single neuron. Open dots are neurons that signaled only timing errors; black dots are neurons that signaled only choice errors; gray dots are neurons that signaled both errors. For each neuron, we computed the mean activity after choice errors and correct responses 100-300 ms afte the primary response in the Fast condition (abscissa), and the mean activity after timing errors and correct responses 100-500 ms after expected reward in the Accurate condition (ordinate).

### Proactive modulation and performance adjustment

Our previous descriptions of visuomotor structures (Heitz and Schall 2012; Reppert et al., 2018) showed that many neurons exhibited proactive modulation of baseline activity before the array appeared, of visual responses, of target selection, and of presaccadic activity. If medial frontal cortex contributes to this modulation, then neurons in SEF must modulate in parallel manner. We tested this.

We measured baseline activity in 40 SEF neurons (20 Da, 20 Eu) with a visual response to the search array, pre-saccadic buildup activity, or a combination thereof. Baseline discharge rate in SEF was significantly elevated under speed constraints in the Fast relative to the Accurate condition (Figures 5A, S4A) (F(1,76) = 12.98, p = 0.0006, BF = 72, two-way ANOVA). We computed z-scored spike counts during the baseline interval 600 ms prior to array appearance and found that the effect of SAT condition was greater during less efficient relative to more efficient search (Figures 5B, S4B) (F(1,76) = 6.01, p = 0.016, BF = 3.46, two-way ANOVA), just as the SAT effect on RT and choice error rate was greater during less efficient relative to more efficient search. Modulation of baseline discharge rate was immediate upon entering the Fast condition (Figures 5C, S4C), as previously found in FEF (Heitz and Schall, 2012) and SC (Reppert et al., 2018). But unlike FEF and SC, single-trial change was asymmetric and of reduced magnitude upon entering the Accurate condition (t_39_ = 2.05, p = 0.047, BF = 1.13).

**Figure 5.**
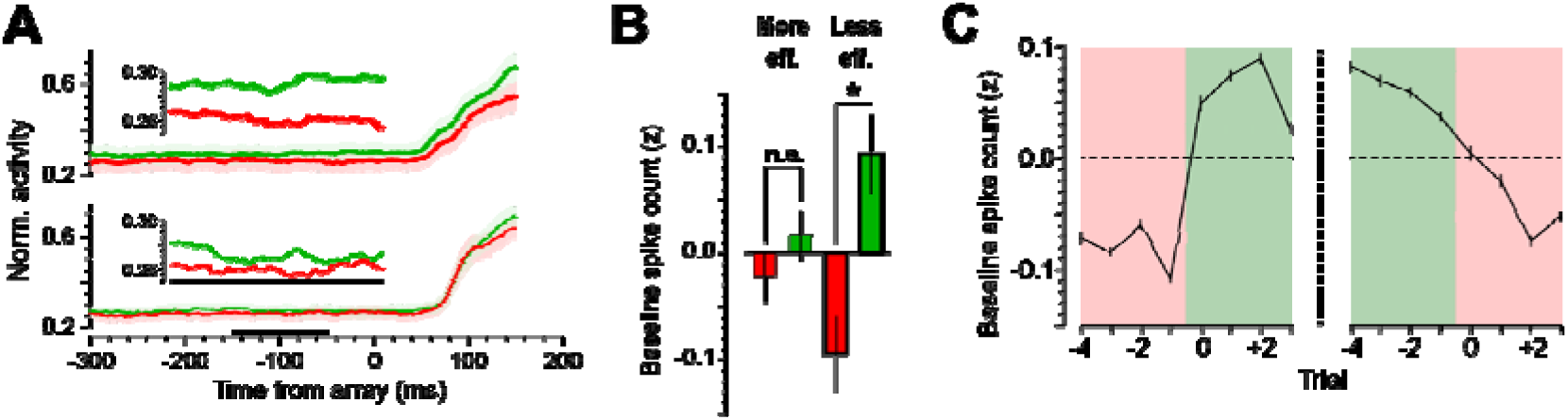
Proactive modulation before array presentation. (**A**) Mean ± SE baseline discharge rate, plotted separately for less efficient (above) and more efficient (below) visual search. Insets show mean discharge rate during the period from 150 ms until 50 ms prior to array appearance (indicated on abscissa). Data averaged across all SEF neurons with either visual response or pre-saccadic buildup activity (n = 40). (**B**) Mean ± SE spike count in the 600-ms interval before array presentation. The effect of SAT condition on baseline activity was greater during less efficient relative to more efficient search, paralleling the elevated effect of SAT on behavior during less efficient relative to more efficient search. (**C**) Change in mean ± SE baseline spike count after SAT cue changes. Unlike SC and FEF, the single-trial modulation of baseline activity was asymmetric and of greater magnitude upon entering the Fast condition than upon entering the Accurate condition.

We observed a visual response to the appearance of the search array in 38 SEF neurons (20 Da, 18 Eu), of which 33 exhibited enhancement (16 Da, 17 Eu), and 5 suppression (4 Da, 1 Eu). Previous work reported elevated visual responsiveness during the Fast condition in FEF (Heitz and Schall, 2012), but not in SC (Reppert et al., 2018). Here we found that SEF visual responsiveness, like baseline activity, was elevated in the Fast relative to the Accurate condition (Figures 6A, S5A) (F(1,62) = 30.2, p = 1.0 × 10^−6^, BF > 1,000, two-way ANOVA). Also as for baseline activity, the SAT effect on visual responsiveness was greater during less efficient relative to more efficient search (Figures 6B, S5B) (F(1,62) = 5.74, p = 0.02, BF = 3.04, two-way ANOVA). Modulation of visual responsiveness was immediate upon a cued change in task condition (Figures 6C, 6D, S5C). Similar to baseline discharge rate, the single-trial change in visual responsiveness was asymmetric. Single-trial modulation was greater upon entering the Fast condition than upon entering the Accurate condition (t_32_ = 3.58, p = 0.001, BF = 29.4).

**Figure 6.**
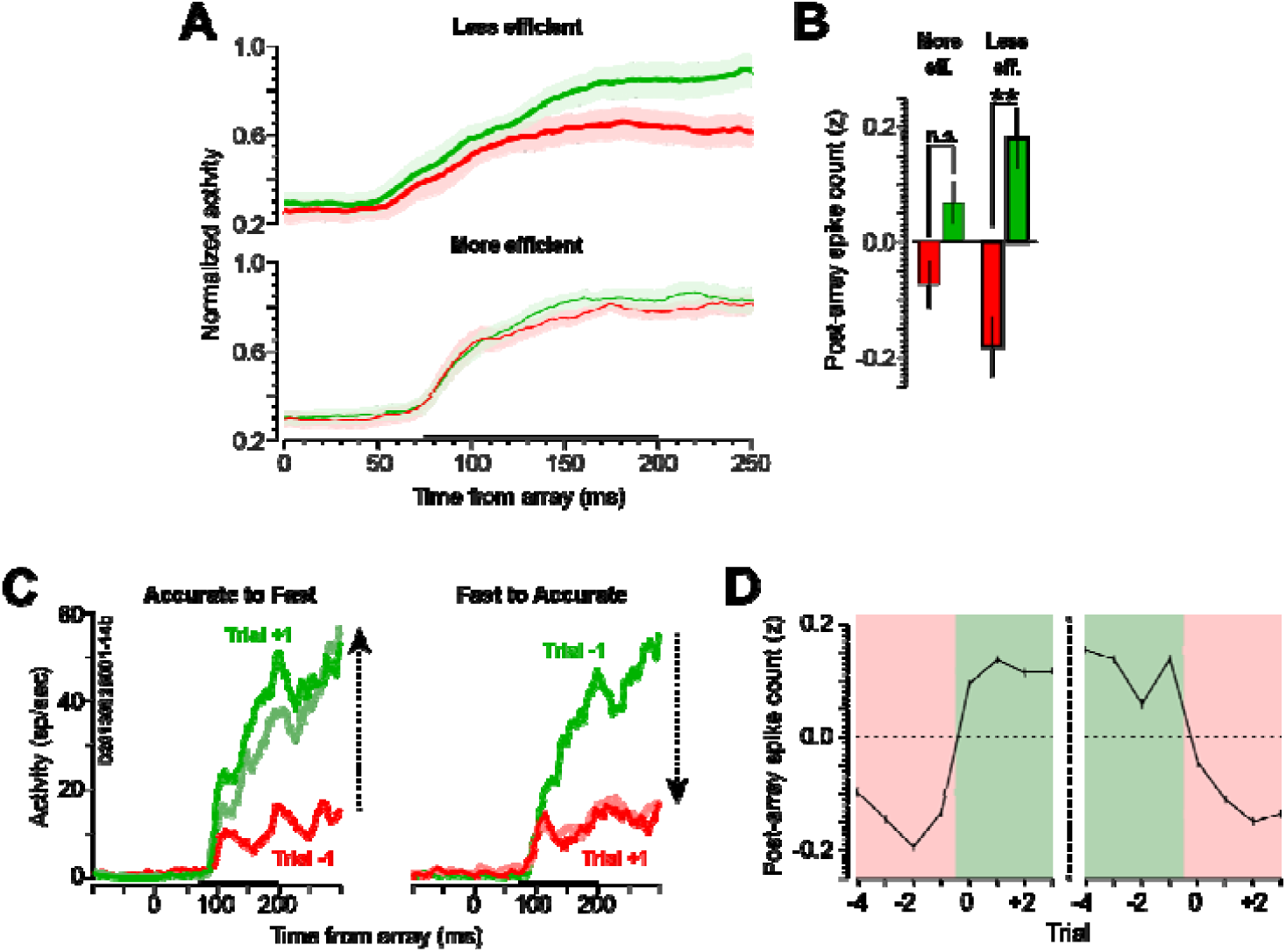
Target salience representation. (**A**) Average SDF when the target appeared in the response field of the neuron during less efficient (above) and more efficient (below) search sessions, combined across common neurons that did not discriminate the target and rare neurons that did. Horizontal bar denotes time window used to quantify visual responsiveness (75-200 ms post-array appearance). (**B**) Mean ± SE spike count during the visual response. The effect of SAT condition on visual responsiveness was greater during less efficient relative to more efficient search. (**C**) SDF of exemplary neuron for triplet of trials preceding (Trial −1), at (Trial 0), and after (Trial +1) transition from Accurate to Fast (left) and vice versa (right). (**D**) Relative to SAT cue changes, mean ± SE visual response discharge rate. Single-trial modulation was greater upon entering the Fast condition than upon entering the Accurate condition.

Overall, the proactive modulation in SEF parallels that observed in visuomotor structures, supporting the notion that medial frontal cortex can be a source of such modulation. However, does such modulation relate to performance adjustments? We observed a different pattern of variation in choice error rate with RT in each task condition (Figure 7A). In the Fast condition, choice error rate decreased with RT (R = −0.22, p = 0.01, BF = 1.72, Pearson correlation), but in the Accurate condition, choice error rate increased with RT (R = 0.41, p = 6.1 × 10^−7^, BF > 1,000), suggesting that monkeys operated on distinct states of responding in two task conditions. Next, for each task condition we assessed the relationship of RT to discharge rate during the baseline and visual response periods and found unexpected patterns of invariance and variation (Figure 7B). Baseline discharge rate was invariant across RT in the Fast condition (R = 0.06, p = 0.38, BF = 0.09), but increased with RT in the Accurate condition (R = 0.27, p = 0.003, BF = 6.16). Visual responsiveness decreased with RT in the Fast condition (R = −0.17, p = 0.010, BF = 1.44), but was invariant across RT in the Accurate condition (R = 0.06, p = 0.38, BF = 0.07).

**Figure 7.**
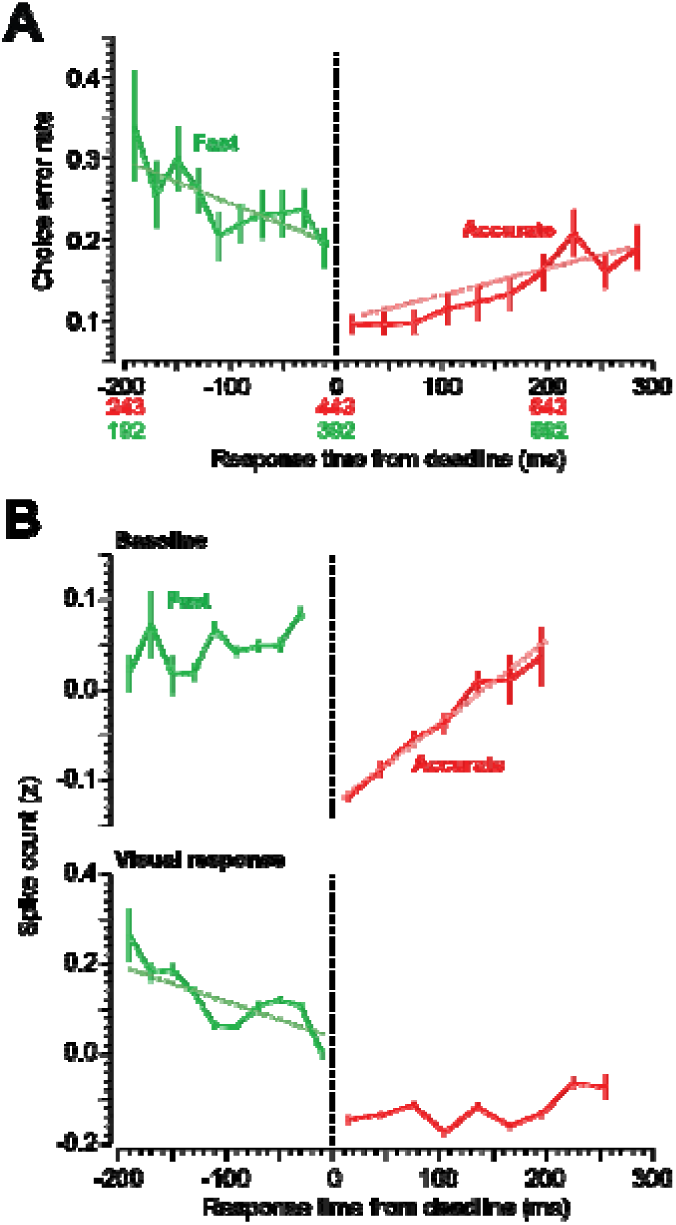
Relationship between neural activity and RT. (**A**) Rate of choice errors vs. RT. The abscissa plots time relative to Fast and Accurate condition deadlines (black text), absolute RT for the Accurate trials (red text), and absolute RT for the Fast trials (green text). Colored dotted lines show best fit to the data. Choice error rate decreased with RT in the Fast condition, but increased with RT in the Accurate condition. (**B**) Discharge rate modulation vs. RT, plotted separately before (above) and after (below) search array presentation. Colored dotted lines show best fit to the data for significant trends. For the baseline period, discharge rate increased prior to prolonged RT in the Accurate but not the Fast condition. For the post-array period, visual response magnitude decreased prior to prolonged RT in the Fast but not the Accurate condition.

## DISCUSSION

Continuing our systematic neurophysiological investigation of SAT in nonhuman primates (Heitz and Schall, 2012, 2013; Reppert et al., 2018; Servant et al., 2019), we now report the first description of single-neuron activity in medial frontal cortex during SAT. We highlight new observations and situate these findings in the context of the neuro-physiological, neuro-imaging, and neuro-computational literatures. We conclude that the collection of new neurophysiological findings across brain structures and laboratories offers insights different from those of cognitive psychology and noninvasive measures. These new insights reveal a more mechanistically plausible account of how the brain accomplishes SAT.

### New insights into SAT of visual search performance

In performing visual search across Fast and Accurate conditions, macaque monkeys produced choice errors (shifting gaze to a distractor) and timing errors (shifting gaze too early or too late). The analysis of the current data offered several new insights. First, we noted incidental but significant differences in visual search efficiency across search arrays (T amongst L’s and L amongst T’s). We found that SAT was exerted to a greater extent during less efficient relative to more efficient search.

As reported previously, error rate and RT adjustments occurred immediately upon SAT condition changes. Here, we noticed asymmetric variation of performance on transitions from Fast to Accurate or from Accurate to Fast conditions, which likely arise from a disposition not to delay responses. The asymmetric variation in the production of saccade across SAT conditions and through trial time will be a crucial test of next-generation models of SAT.

Following choice errors, we noticed that monkeys commonly produced a second saccade to the location where the target had been. Neural correlates of this will be discussed below. Curiously, the interval between the choice error and corrective saccades was significantly longer during the Accurate relative to the Fast condition. This indicates that SAT is a prolonged state that persists beyond the formal trial (Sedaghat-Nejad et al., 2019). This is an additional feature that next-generation response accumulator models must account for.

### New insights into performance monitoring with SAT

The visual search task allowed us to dissociate errors of choice (shifting gaze to a distractor) from errors of timing (shifting gaze too early or too late according to SAT cues). This dissociation provided new information about performance monitoring in medial frontal cortex. As reported previously (Abzug and Sommer, 2018; Nakamura et al., 2005; Purcell et al., 2012; Sajad et al., 2019; Stuphorn et al., 2000), SEF neurons signaled when monkeys made choice errors. Following choice errors, even though the search array was extinguished, monkeys commonly made corrective saccades to the location where the target had been. Saccades correcting fixation errors in visual search are common in monkeys (Murthy et al., 2007).

These choice error corrections afforded an opportunity to investigate relationships between error signaling and error adjustment, which have been difficult to observe (Cohen and Ranganath, 2007; Dudschig and Jentzsch, 2009; Fiehler et al., 2005; Gehring and Fencsik, 2001; Gehring et al., 2012; Hajcak et al., 2003) (but see Sajad et al., 2019). We found that larger error-related modulation was associated with a greater likelihood of production of corrective saccades. Such a relationship has not been reported to our knowledge. Moreover, we also found that the corrective saccades were produced ∼ 200 ms after neurons signaled the choice error. This contrasts sharply with our observation during a dynamic visual search task of corrective saccades produced faster than error signals (Murthy et al. 2007). While the differences are surely just a consequence of different task conditions, they highlight the flexible nature of behavior.

Following timing errors, the search array remained visible until the time of reward delivery. Replicating earlier studies in medial frontal cortex by ourselves (Ito et al. 2003) and others (Roesch and Olson, 2003; Seo and Lee, 2009; So and Stuphorn, 2010), neurons signaled when monkeys did not receive expected juice reward. This modulation has been described as reward prediction error (Schultz, 2016), or surprise (Kawaguchi et al., 2015). We found that overt errors and reward prediction errors were signaled independently by some neurons, and collectively by others. The diversity of functional types of neurons in medial frontal cortex emphasizes that unitary hypotheses about medial frontal function are oversimplified and highlights the need for next generation accumulator models that incorporate and explain the utility of this diversity. In general, the results corroborate the contribution of SEF to performance monitoring and reveal additional properties of medial frontal cortex circuitry.

### New insights into proactive state changes during SAT

As reported in FEF (Heitz and Schall, 2012) and SC (Reppert et al., 2018), neurons in SEF produced higher discharge rates in the Fast condition during the baseline period and also in response to the array. Higher discharge rates in Fast relative to Accurate SAT trials was also reported in LIP (Hanks et al.2014), in skeletal motor cortical areas (Thura & Cisek 2016) and in the basal ganglia (Thura and Cisek 2017). These remarkably consistent observations across structures and laboratories demonstrates that this proactive state change is broadcast broadly in cortical and subcortical structures. We discovered that this modulation in SEF varied with visual search efficiency, i.e., task difficulty. In general, the proactive modulation was greater when search efficiency was lower (task difficulty was higher). This finding parallels many other studies that have reported that activation in medial frontal cortex increases with task difficulty and error rate, and it reveals another, unexpected layer of complexity in the neural mechanisms of SAT.

As reported previously (Purcell et al. 2012), and unlike in FEF and SC, few neurons in SEF contributed to the process of saccade target selection. Among the few neurons, though, this target selection process occupied the same amount of time as that measured in FEF and SC of the same monkeys and exhibited the same delay in the Accurate relative to the Fast conditions. The simultaneity of target-distractor modulation and parallel modulation across SAT conditions across regions are consistent with an interactive network or common source. One possibility is that SEF receives target selection signals from FEF directly and from SC indirectly via the thalamus. However, such connectivity is unlikely to offer a complete explanation of this process in SEF. Relative to the previous study, though, we found a higher fraction of SEF neurons signaling search target location (24% compared to 2%). This difference can be appreciated based on differences in the task conditions. The previous study sampled in monkeys performing a highly efficient, color singleton (pop-out) search, while the present study sampled in monkeys performing a less efficient search for form (T/L) under different SAT instructions. While further research is needed to determine what factors explain the difference, the overall pattern of these findings is consistent with previous work showing that when tasks are more demanding, medial frontal cortex contributes more resources.

### New insights into mechanisms of SAT

When viewed from the perspectives of formal models of SAT (Bogacz et al., 2010) and previous studies of SAT using noninvasive measures of physiology (Forstmann et al., 2008, 2010; Ivanoff et al., 2008; van Maanen et al., 2011; Murphy et al., 2016; Steinemann et al., 2018; van Veen et al., 2008; Weigard et al., 2018; Wenzlaff et al., 2011), coupled with prior extensive knowledge about the connectivity and functional properties of the brain circuits producing visually guided saccades (e.g., Liversedge et al., 2011), and previous studies of FEF (Heitz and Schall, 2012; Servant et al., 2019) and SC (Reppert et al., 2018), LIP (Hanks et al., 2014), skeletomotor cortex (Thura and Cisek, 2016) and basal ganglia (Thura and Cisek, 2017), the current findings offer several new insights into the neural mechanisms of SAT.

First, as noted previously (Heitz and Schall, 2012, 2013), formal models of SAT in terms of simple stochastic accumulation of evidence (Bogacz et al., 2010) do not explain all of the processes changing in the brain (cf. Cassey et al., 2014). This does not mean that the psychological process models are no longer a useful explanation of behavior; it simply means they are not transparent explanations of the brain. An implication of this lack of transparency will be highlighted below. The new observations presented here of diverse modulation in SEF associated with correct and error performance demonstrate that a more complete formal model should incorporate additional elements like error monitoring, reward monitoring, and pro-active preparation.

Second, although the contribution of medial frontal cortex (as well as other structures) to SAT adjustments is clear from non-invasive human studies, these new neurophysiological results seem consistent with some observations but raise questions about previous interpretations derived from non-invasive measures. For example, the robust neurophysiological finding of elevated discharge rates at baseline is consistent with the pre-stimulus preparatory changes reported in neuroimaging studies of SAT (e.g., Forstmann et al., 2008; van Veen et al., 2008).

However, the consistently larger response to task stimuli and during responses in Fast relative to Accurate trials is not consistent with neuroimaging reports of reduced transient response-related activation during speed emphasis (Ivanoff et al., 2008; van Veen et al., 2008). In addition, some neuroimaging studies conclude that emphasis on speed or accuracy entail different networks or network configurations (e.g., Forstmann et al., 2008; Weigard et al., 2018). This conclusion is difficult to reconcile with the neurophysiological data from this and the other studies showing only quantitative and not qualitative differences across SAT conditions. We note that comparing neurophysiological and neuroimaging results entails assumptions about the mapping of sampled spike rates and BOLD activation that cannot be taken for granted (Logothetis and Wandell, 2004).

Meanwhile, evidence from studies of SAT measuring event-related potentials has led to the general conclusion that SAT operates on response preparation, but not earlier operations such as salience representation (van der Lubbe et al., 2001; Osman et al., 2000; Rinkenauer et al., 2004). However, here we found pronounced modulation across SAT conditions observed before and in response to array presentation. Studies of the effects of SAT on error-related brain activity have reported enhanced error signals (both Ne/ERN and Pe) during trials emphasizing accuracy over speed (e.g., Arbel and Donchin, 2009; Gehring et al., 1993). A more recent ERP study investigated error monitoring during SAT (Steinhauser and Yeung, 2012) to test predictions derived from a simulation that used response criterion shifts to accomplish SAT (Steinhauser et al., 2008). The average amplitude of the error-related negativity was not significantly different across SAT conditions, but the post-error positivity was reduced in the Accurate relative to the Fast trials, and the trial-by-trial amplitude of the positivity predicted whether participants would report errors. Our neurophysiological findings are inconsistent with variation of Ne/ERN and Pe across SAT conditions, but consistent with the relation of post-error positivity and error awareness (if corrective saccades assay awareness). The disparity between previous ERP and current neurophysiological findings may be due to flawed assumptions about the mechanisms of SAT. If response threshold does not actually change across task conditions, then this has implications for the motivation, execution, and interpretation of previous studies of SAT. In particular, quantitative predictions derived from a simulation that seeks to investigate theories of error detection based on this standard assumption of SAT (e.g., Steinhauser et al., 2008) may be questionable. Differences between the EEG findings and the neurophysiological findings are surely also related to the currently uncertain relationship between EEG and neural spiking. Research on this problem is ongoing (Sajad et al., 2019). Alternatively, the simulation leading to the prediction may be incorrect because the brain does not actually change response threshold as assumed by the standard SAT models.

Finally, a prominent hypothesis suggests that SAT is accomplished through circuits linking medial frontal lobe to the basal ganglia (e.g., Forstmann et al. 2008, 2010). Given the established connectivity of SEF (and FEF) with dorsal striatum (Parthasarathy et al., 1992), our findings are relevant for evaluating aspects of that hypothesis. Indeed, the proactive modulation of activity across SAT conditions that we discovered in FEF (Heitz and Schall, 2012) and have also found here in SEF is also present in the basal ganglia (Thura and Cisek, 2017). Hence, our new findings, taken in conjunction with other neurophysiological and neuroanatomical findings, support a new mechanistic perspective on neural mechanisms of SAT. For example, previous work emphasized a contribution of the hyper-direct pathway from medial frontal cortex to the subthalamic nucleus (Forstmann et al., 2010; Frank, 2006). However, anatomical studies have confirmed that SEF does not project to the subthalamic nucleus (Huerta and Kaas 1990).

Although differences of effector and of species must be considered, we believe the bulk of evidence indicates that medial frontal areas play more of a supervisory role, akin to a critic of the visuo-motor circuit actor. Previous research has shown that SEF, similar to preSMA (Scangos and Stuphorn, 2010), does not directly control the timing of movements (Stuphorn et al., 2010). Instead, SEF, like preSMA, monitors performance and exerts indirect influence on motor processes to adjust RT (Scangos et al., 2013; Stuphorn et al., 2010). Such indirect influence can be described as urgency, as suggested by others (e.g., Thura and Cisek, 2016), and implemented in abstract spiking network models (Lo et al., 2015). Earlier research showed that intracortical electrical microstimulation of sites in SEF can speed or slow RT according to task context (Stuphorn and Schall, 2006). Also, previous research has demonstrated that saccades are initiated when the discharge rates of accumulation neurons in FEF and SC reach a threshold (e.g., Hanes and Schall, 1996). The activity of these neurons corresponds precisely with the evidence accumulation process supporting visual search performance (Purcell et al., 2010, 2012).

Another neurophysiological study investigated how strategic changes of RT are implemented in FEF and SC (Pouget et al., 2011). RT was advanced or delayed not by changes in response threshold, but instead by changes in the time of accumulation initiation. Thus, in contrast to previous suggestions about medial frontal contributions to the evidence accumulation process (e.g., Forstmann et al., 2008; Ivanoff et al., 2008), SEF does not influence the excursion of the accumulation process but rather modifies the time when accumulation starts. A neuro-computational model of perceptual decision making has demonstrated that variation of onset of accumulation can arise by adjusting a gate on the level of evidence necessary to begin accumulation (Purcell et al., 2012). The gated accumulator model was applied to neurophysiological data obtained in FEF of three monkeys that performed visual search under SAT (Servant et al., 2019). The model fits exposed differences in SAT strategies across monkeys and highlighted the actual complexity of the neural mechanisms of SAT.

Models of SAT of perceptual decision-making to date have accounted for the systematic adjustments of performance through changes in the parameters governing evidence accumulation. None of the models actually explain how those parameters are made to change. Many believe that medial frontal cortex exerts executive control over sensory-motor circuits. The results of this first neurophysiological investigation of medial frontal cortex during speed-accuracy tradeoff demonstrate the diversity of signals and modulations required to make the parameter changes needed to accomplish SAT.

## METHODS AND MATERIALS

### Subjects

Two bonnet macaques (*M. radiata*), identified as Da and Eu, performed the visual search task. All surgical and experimental procedures were completed in accordance with the National Institutes of Health Guide for the Care and Use of Laboratory Animals, and were approved by the Vanderbilt Institutional Animal Care and Use Committee in accordance with the United States Department of Agriculture and Public Health Service Policy on Humane Care and Use of Laboratory Animals.

### Task

Monkeys performed a form (T/L) visual search task for a target item presented amongst seven iso-eccentric distractor (L/T) items. Trials began when monkeys fixated gaze on a central cue for ∼ 1 sec. Monkeys were extensively trained to associate the color of the fixation cue (red or green) with the SAT task condition (Accurate or Fast). After fixation of the central cue, the visual search array of form stimuli appeared, of which one form was the target for that session (T or L). All stimuli (both target and distractors) were oriented randomly in cardinal directions. During several sessions, all distractor items were oriented identically. Monkeys Da and Eu completed 8 and 6 sessions with 894 ± 76 (Da) and 837 ± 74 (Eu) correct trials per session, respectively.

Trials were run in blocks lasting 5 or 10 trials, alternating between Fast and Accurate conditions. In the Accurate condition, saccades to the target form were rewarded if the RT exceeded an unsignaled deadline (Da: 434 ± 7 ms, Eu: 454 ± 10 ms). Following correct responses, monkeys fixated the target form for ∼ 700 ms, until the fixation cue and array were extinguished and fluid reward delivered. Saccades directed to a distractor (choice errors) and saccades that were executed too early (timing errors) were followed by a 4 sec time out. No explicit feedback was given after timing errors. In the Fast condition, saccades to the target form were rewarded if the RT preceded the unsignaled deadline (Da: 362 ±17 ms, Eu: 433 ± 26 ms). Responses executed after the deadline were followed by a 4 sec time out. Saccades directed to a distractor executed before the deadline had no time out. Monkeys had difficulty discriminating lack of reward from choice errors and timing errors. Therefore, the display was removed on 25%-50% of timing error trials during the Fast condition. Monkeys thus learned that reinforcement was only available prior to the deadline.

### Behavioral data acquisition and analysis

All gaze data were collected with video-oculography (Eyelink 1000, SR Research, Kanata, Ontario, Canada) and analyzed offline using Matlab R2019a (The MathWorks, Inc.). Gaze position data were filtered offline with a 3^rd^-order Butterworth low-pass filter (cutoff frequency = 80 Hz). The data were then differentiated to velocity traces, and a cutoff velocity of 30°/sec was used to identify all saccades. Response time was determined as the first time-point when the velocity trace of the task-relevant saccade crossed the 30°/sec threshold.

Responses were deemed timing errors if they were initiated after the unsignaled deadline in the Fast condition, or before the deadline in the Accurate condition. Responses were deemed choice errors if the saccade endpoint did not fall within the angular octant occupied by the target form T/L.

We assessed change in behavioral parameters such as RT, timing error rate, and choice error rate across trials. For this analysis, we identified all trials with a switch from Fast to Accurate condition, or vice versa, and aligned trials accordingly. We then computed mean values across all sessions.

### Neural data acquisition

Details of the methods have been reported previously (Heitz and Schall, 2012; Reppert et al., 2018). During recordings, monkeys sat in enclosed primate chairs with heads restrained in front of a CRT monitor. Daily recording protocols were consistent across monkeys and sessions. After advancing the recording electrode to the desired depth, 3-4 h elapsed to allow stabilized recordings. This waiting period resulted in consistently stable recordings.

Intracranial data were recorded from the SEF using tungsten microelectrodes (2-4 MΩ, FHC, Inc.). Location was verified by evoking eye movements via low-threshold (<50 μA) microstimulation. The number of electrodes lowered during a single session ranged from one to nine, including the use of dual FHC (1×4 or 2×2) manifolds. Neurons were sampled in SEF and also in FEF in one monkey and in SC in the other. Microelectrodes were referenced to a guide tube in contact with the dura mater. Single-unit waveforms were sampled at 40 kHz, isolated online, and resorted offline (Offline Sorter, Plexon Inc., Dallas, TX, USA). Spike trains were convolved with a kernel that resembled a postsynaptic potential to create a spike density function (SDF) (τ_growth_ = 1 ms, τ_decay_ = 20 ms).

### Neuron classification

We characterized SEF neurons as having visually-responsive activity, saccade-related activity, and/or error-related activity during the search task. We classified neurons as visually-responsive based on examination of discharge rate after appearance of the target stimulus during a memory-guided saccade task. We first determined all neurons with modulation of activity for at least one of the eight possible stimulus locations during the interval from 0 to 400 ms post-stimulus presentation. Visually-responsive neurons were classified as either stimulus-enhanced (n = 16 Da, n = 17 Eu) or stimulus-suppressed (4 Da, 1 Eu), depending on the direction of modulation relative to the baseline discharge rate of the neuron. We determined the response field of all visually-responsive neurons as the sum of all directions for which stimulus presentation evoked a visual response during the memory-guided task. We classified neurons as saccade-related based on examination of discharge rate before execution of the saccadic response during a memory-guided saccade task (n = 12 Da, n = 10 Eu). We determined the movement field of all saccade-related neurons as the sum of all directions for which pre-saccade activity was significantly elevated during this task.

### Normalization of spiking activity

We normalized the activity of neurons on an epoch-specific basis. Baseline activity was normalized to the mean activity across all trials (both task conditions) during the 700-ms period prior to array appearance. Post-stimulus appearance activity was normalized to the maximum of the mean SDF in the Fast condition during the 400 ms window immediately after the appearance of the array stimulus. Peri-saccade activity was normalized to the maximum of the mean SDF in the Fast condition during the window from 100 ms pre- to 100 ms-post primary saccadic response.

### Analyses of proactive modulation and visual response

To test for significant effects of SAT condition and search efficiency on baseline activity (interval [-600 ms, +20 ms] from array appearance), we used two-way between subjects ANOVA with factors condition (Fast / Accurate) and search efficiency (more / less). We first computed single-trial spike counts in the baseline interval and converted these to z-scored counts on an individual neuron basis. To test for single-trial change in baseline activity, we computed the difference in z-scored spike counts for the last trial before and first trial after a change in task condition. We used a paired t-test to determine whether single-trial change was greater upon entering the Fast or the Accurate SAT condition.

We computed the visual response latency as a function of the baseline discharge rate. Specifically, we calcualated the mean (μ) and standard deviation (σ) of the SDF during the baseline period, and then compared the past-array activity to these values. We determined the first time at which visual response activity reached 6 SD greater than mean baseline activity and remained above this value for at least 50 ms. Given that latency values were similar, we averaged values across Fast and Accurate SAT conditions. We computed the magnitude of the visual response to the search array as the mean of the baseline-subtracted spike count during the time window [+75 ms, +200 ms] from array appearance. As for baseline discharge rate, we used two-way between-subjects ANOVA to test for main effects of task condition and search efficiency on magnitude of the visual response.

### Analysis of target selection

To test for the presence of target selection during the visual response, we assessed single-trial spike density functions during the 350 ms interval immediately post-visual response onset. We performed a one-sided Mann-Whitney U-test on a sample-by-sample basis to determine the first timepoint for which the null hypothesis of equal discharge rate for trials with target in RF vs. distractor in RF was rejected (α < 0.01) for a minimum duration of 50 ms. We labeled this timepoint the time of target selection. In few instances, when this algorithm failed to detect appropriate TST, we adjusted our estimate based upon visual inspection of the SDF (i.e., the differences between SDF with target in the neuron’s RF vs. distractor in the RF).

### Analysis of post-response choice error-related activity

To test for the presence of choice error-related activity, we compared post-primary response activity on correct and error trials during the search task. We identified 35 units (Da: 20, Eu: 13) with facilitation or suppression of activity on choice error trials.

Given that a secondary saccade was generated ∼ 300 ms after the error (Figure S2C), it was possible that neurons that we interpreted as error-related were modulated by generation of another saccade. To control for the confound of movement-related activity, we split choice error trials into two groups split on the median latency of the secondary saccade. We quantified the onset time of choice error-related modulation separately for each group. We used the following statistic to determine neurons that responded to generation of the secondary saccade: 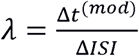 where *t*^*(mod)*^ is the time of neural modulation and *ISI* is inter-saccade interval between the end of the primary saccade and initiation of the second saccade. We removed 4 neurons for which *λ* > 0.5 from further error-related analyses, leaving 28 error-related neurons (Figures S2F, S2G).

### Time of choice- and timing-error related modulation

To determine when neurons encoded for errors, we computed ms-by-ms Wilcoxon rank-sum tests, evaluating the null hypothesis that there was no difference between activity on correct and on error trials. For choice errors, we computed these tests on the interval from −200 ms to 800 ms from primary response initiation. For timing errors, we computed these tests on the interval from −200 ms to 800 ms from time of expected reward. For both trial types, time of onset of error encoding was the first successive 100 ms with a significant difference at the p < 0.05 level.

### Statistical analyses

For measures of task parameters (e.g., response deadline) and behavioral measures, we report mean ± standard error of the mean across all recording sessions. All t-tests are paired, unless otherwise stated. To test for main effects of task condition (Fast / Accurate) and search efficiency (more / less) on measures of behavior and neurophysiology, we employed two-way between-subjects ANOVA. We report values for Bayes Factor alongside all t-statistics and F-statistics, where appropriate. All values for t-statistics, F-statistics, and BF were computed using the R programming language (GNU General Public License, Free Software Foundation, Inc.).

## Acknowledgments

This work was supported by F32-EY028846 to T.R.R., F32-EY019851 to R.P.H., T32-EY007135 for T.R.R., and R01-EY08890, P30-EY08126, U54-HD083211 and by Robin and Richard Patton through the E. Bronson Ingram Chair in Neuroscience. We thank J. Elsey, M. Feurtado, M. Maddox, S. Motorny, J. Parker, M. Schall, and L. Toy for animal care and other technical assistance. We thank A. Sajad for helpful advice and guidance.

## Conflict of Interest

The authors declare no competing financial interests.

