## Supplemental information for "Neural Mechanisms for Executive Control of Speed-Accuracy Tradeoff"

**Supplemental Results**

*Visual response: Target selection*

Previous work reported very few SEF neurons contributing to target discrimination during visual search (Purcell et al., 2012). Here we found target selection in the activity of 8 of 33 visually-responsive SEF neurons. Although a small sample, the time of target selection was delayed in the Accurate relative to the Fast condition during more efficient but not less efficient search (F(1,6) = 7.19, p = 0.036, BF = 2.48, two-way ANOVA interaction effect). This timing difference parallels observations in FEF and SC (Heitz and Schall, 2012; Reppert et al., 2018).

**Supplemental Figures and Legends**

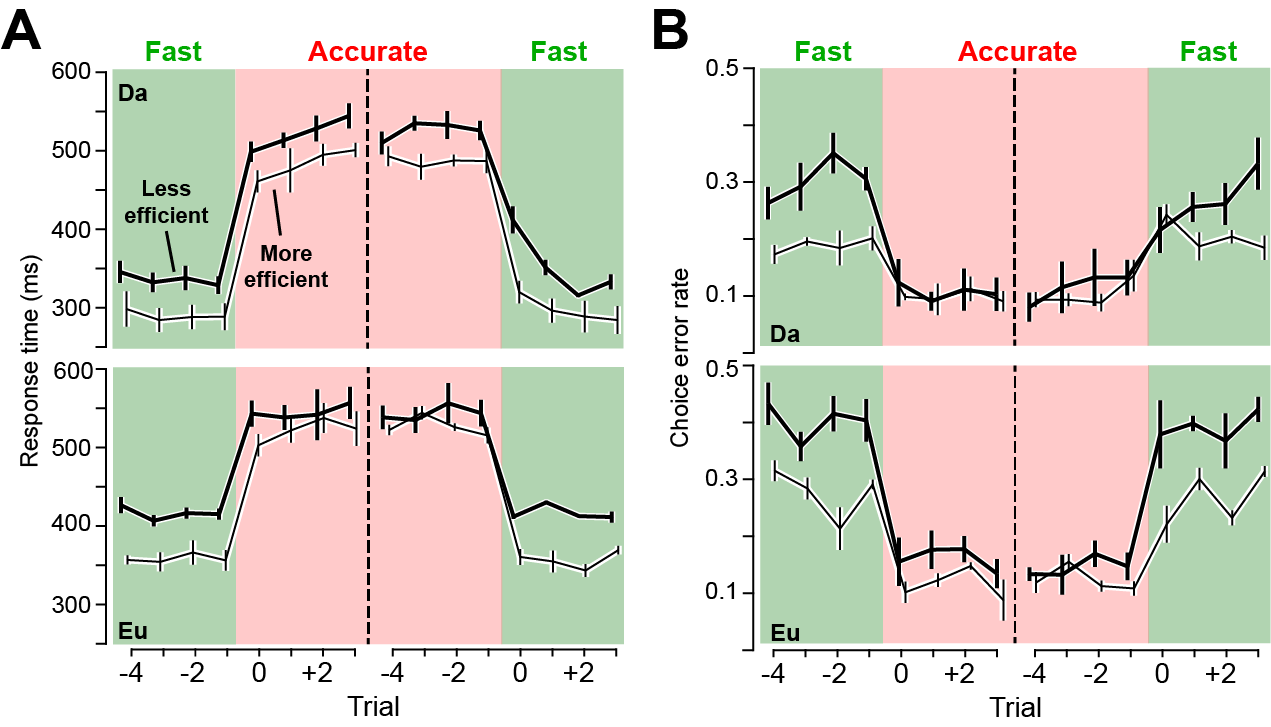

**Figure S1.** Trial-to-trial change in speed-accuracy tradeoff of visual search.

(**A**) Change in mean ± SE RT after SAT cue changes. Monkeys adjusted RT immediately. Data plotted separately for more efficient (thin lines) and less efficient (thick lines) visual search, for monkeys Da (top) and Eu (bottom).

(**B**) Change in mean ± SE choice error rate after SAT cue changes. Error rate changed immediately.

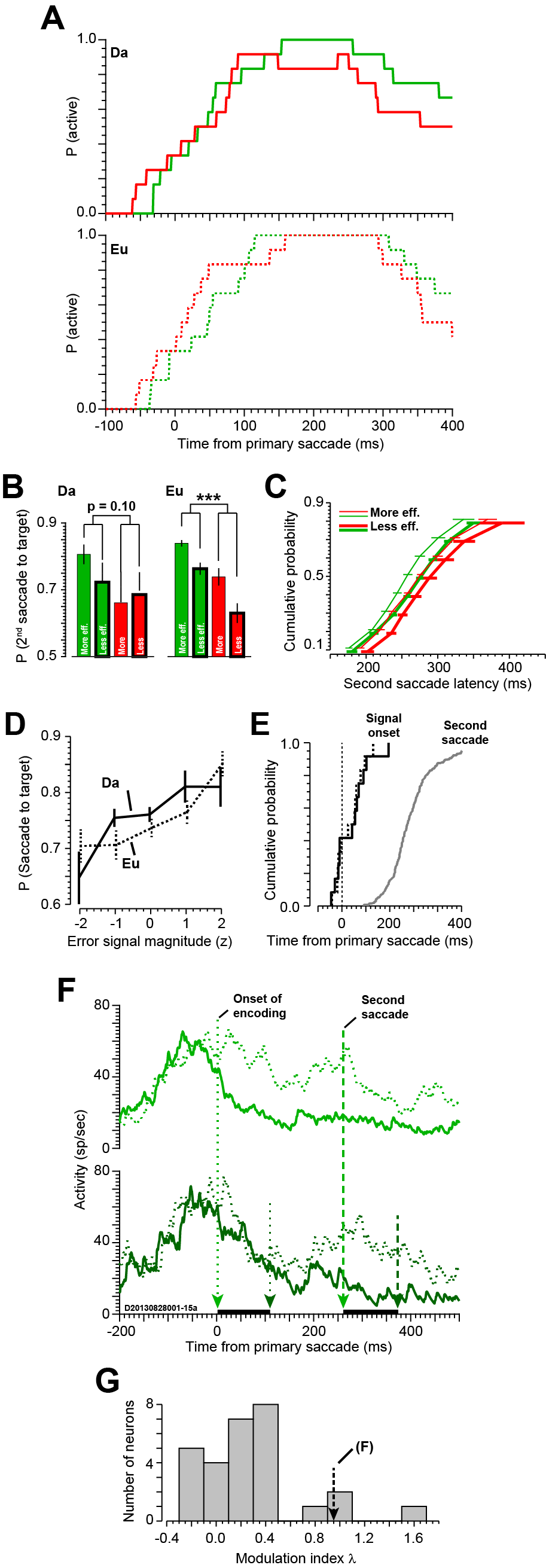

**Figure S2**. Signaling choice errors and correction.

(**A**) Probability density of choice error-related modulation across the sample, plotted separately for monkeys Da (above) and Eu (beneath).

(**B**) Mean ± SE probability of second saccade to target.

(**C**) Cumulative probability ± SE of latency of the second saccade, split by task condition (Fast – green, Accurate – red) and search efficiency (more efficient – thin lines, less efficient – thick lines).

(**D**) Mean ± SE probability of second saccade directed to the target as a function of quantiles of z-scored error-related activity across neurons. For both monkeys, the likelihood of a corrective saccade to the target increased with magnitude of the error signal.

(**E**) Cumulative distributions of time of choice error-related modulation and time of second saccade initiation, plotted separately for monkeys Da (solid line) and Eu (dashed line). Error-related modulation preceded the second saccade by ~200 ms on average.

(**F**) Single-neuron example of the procedure used to control for modulation associated with execution of the second saccade. Average SDF for median-split groups of Fast condition trials with short ISI (top) and long ISI (bottom) are shown in light and dark green. Dark lines on abscissa represent temporal shifts of neural modulation and time of second saccade. For this neuron, initiation of the error signal was tightly linked to the time of the second saccade. We removed all such neurons from classification as choice error-modulated.

(**G**) Histogram of index used to control for modulation associated with the second saccade. Arrow is value associated with example neuron in (F).

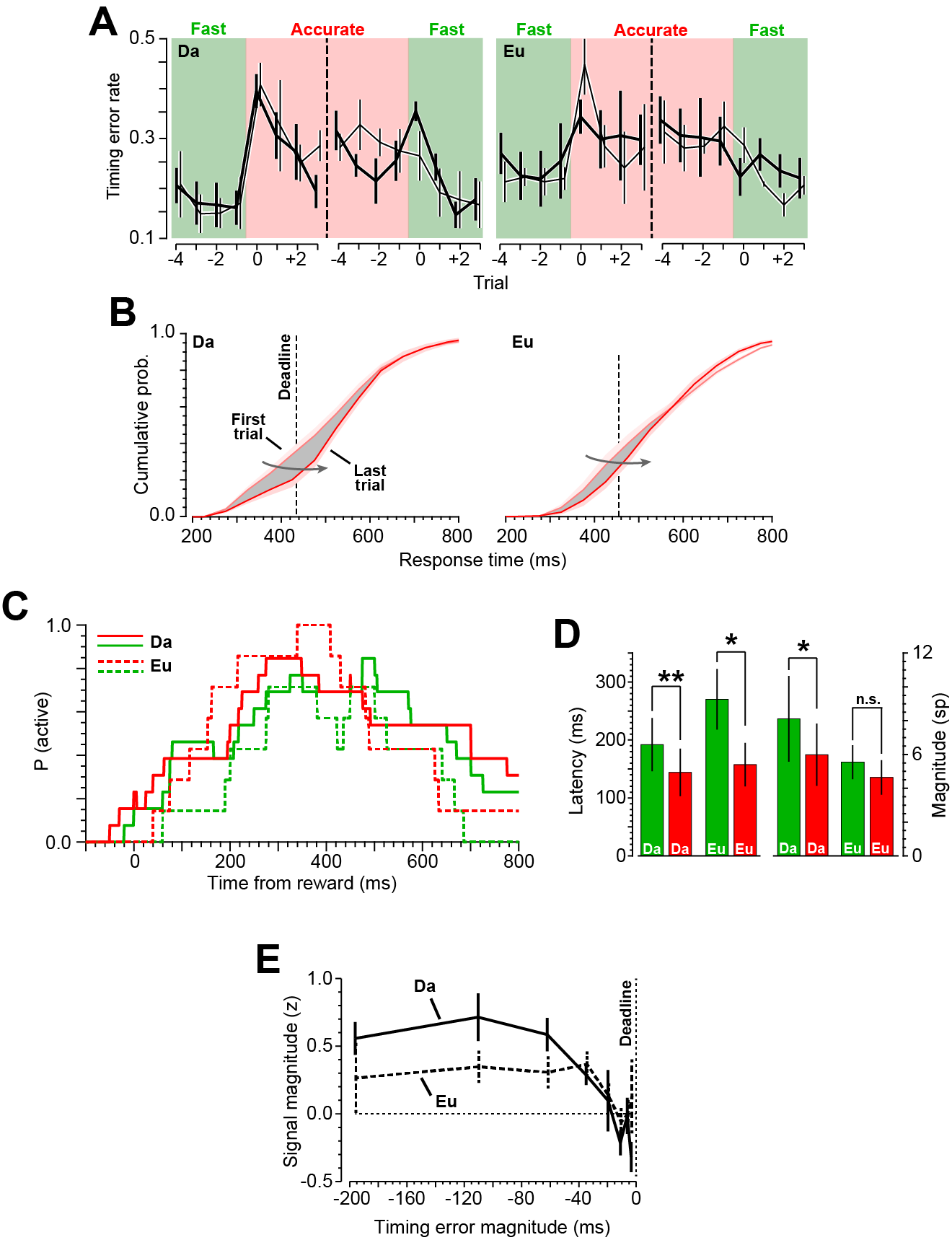

**Figure S3.** Signaling reward prediction error after timing errors.

(**A**) Mean ± SE rate of saccade timing errors relative to cued change in task condition. In both monkeys, changes in timing error rate were immediate upon a cued change in task condition.

(**B**) Cumulative distributions of RT across the first and last trials of the Accurate condition blocks.

(**C**) Probability density of reward prediction error signaling relative to time of expected reward, plotted separately for monkeys Da (solid lines) and Eu (dashed lines).

(**D**) Mean ± SE latency and magnitude of the RPE signal, shown separately for the two monkeys.

(**E**) Magnitude of z-scored error signal as a function of the magnitude of the timing error relative to the SAT deadline (indicated).

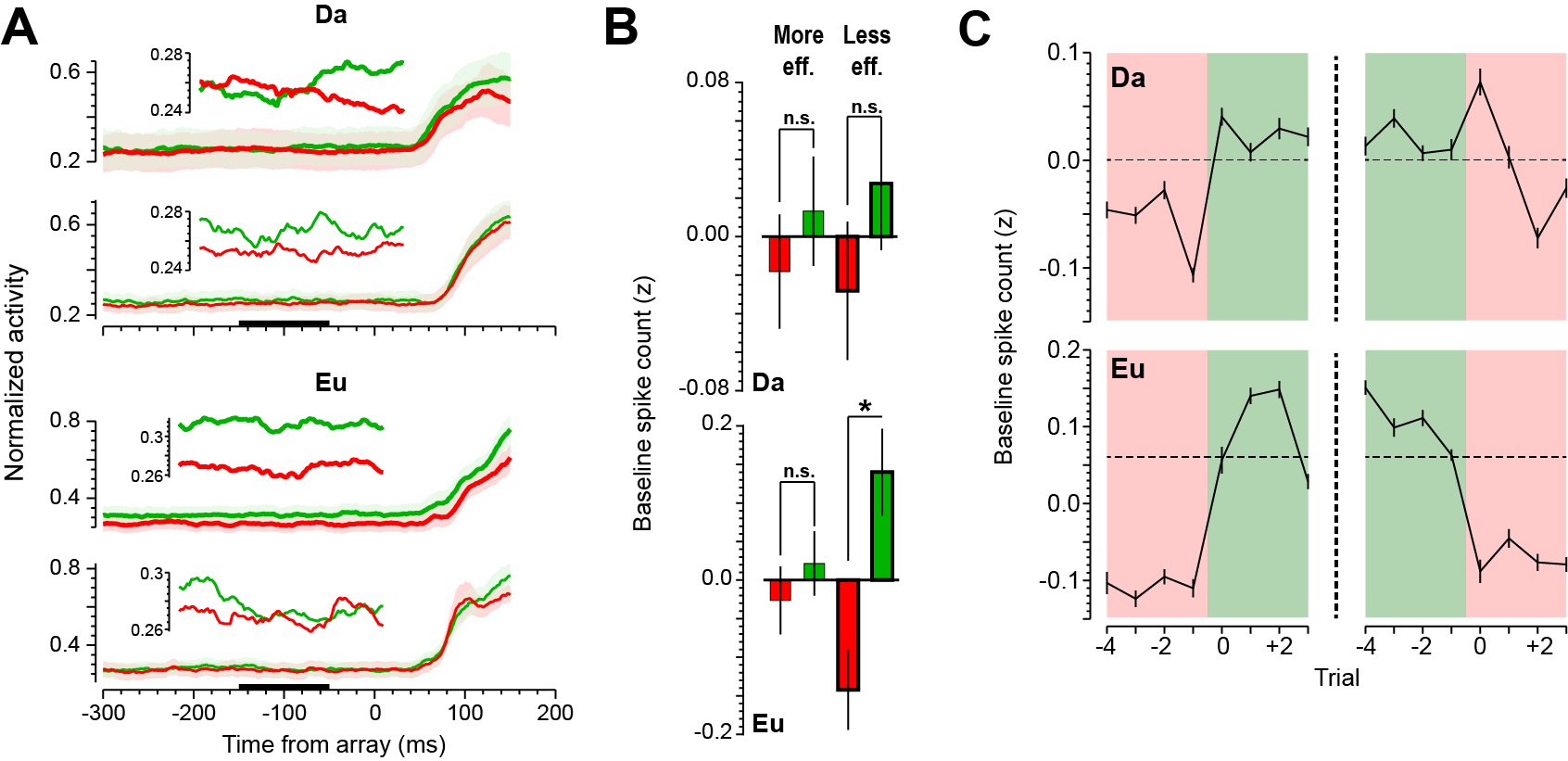

**Figure S4.** Proactive modulation in SEF before array presentation.

(**A**) Mean ± SE baseline discharge rate, plotted separately for less efficient (above) and more efficient (below) visual search. Insets show mean discharge rate during the period from 150 ms until 50 ms prior to array appearance. Data plotted separately for monkeys Da (top) and Eu (bottom).

(**B**) Mean ± SE z-scored spike count in the 600-ms interval before array presentation. The effect of SAT condition on baseline activity was greater during less efficient relative to more efficient search.

(**C**) Change in mean ± SE z-scored baseline spike count after SAT cue changes. In both monkeys, change in baseline activity was immediate upon a cued change in task condition, with greater single-trial modulation upon entering the Fast relative to the Accurate condition.

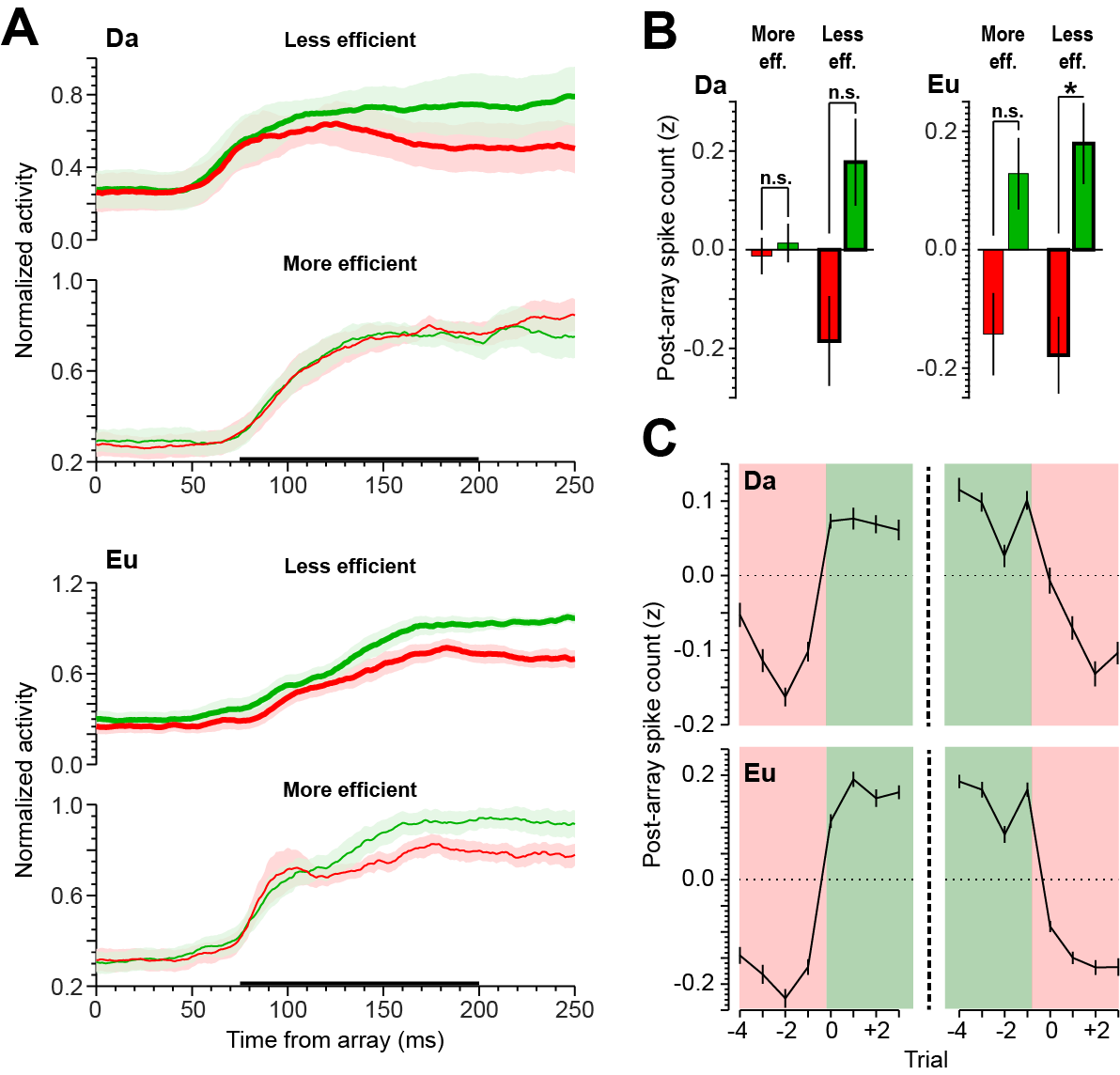

**Figure S5.** Target salience representation.

(**A**) Average SDF when the target appeared in the response field of the neuron during less efficient (above) and more efficient (below) search sessions, combined across common neurons that did not discriminate the target and rare neurons that did. Horizontal bar denotes time window used to quantify visual responsiveness (75-200 ms post-array appearance). Data plotted separately for monkeys Da (top) and Eu (bottom).

(**B**) Mean ± SE z-scored spike count during the visual response.

(**C**) Mean ± SE change in z-scored visual response spike count after cued condition switch. In both monkeys, single-trial modulation was greater upon entering the Fast condition than upon entering the Accurate condition.
